# Generation and timing of graded responses to morphogen gradients

**DOI:** 10.1101/2021.05.11.443662

**Authors:** Shari Carmon, Felix Jonas, Naama Barkai, Eyal D. Schejter, Ben-Zion Shilo

## Abstract

Morphogen gradients are known to subdivide a naïve cell field into distinct zones of gene expression. Here we examine whether morphogens can also induce a graded response within such domains. To this end we explore the role of the Dorsal protein nuclear gradient along the dorso-ventral axis in defining the graded pattern of actomyosin constriction that initiates gastrulation in early *Drosophila* embryos. Two complementary mechanisms for graded accumulation of mRNAs of critical zygotic target genes were identified. First, activation of target-gene expression expands over time from the ventral-most region of high nuclear Dorsal to lateral regions where the levels are lower, due to a Dorsal-dependent priming probability of transcription sites. Thus, sites that are activated earlier will lead to more mRNA accumulation. Second, once the sites are primed, the rate of Pol II loading is also dependent on Dorsal levels. Morphological restrictions require that translation of the graded mRNA be delayed until completion of embryonic cell formation. Such timing is achieved by large introns, that provide a delay in production of the mature mRNAs.

## Introduction

Morphogen gradients manifest positional information by graded activation of signaling pathways that culminate in activation of transcription factors. This activation leads to the induction of target genes that establish the fate of the tissue. The most common outcome of morphogen gradients is the division of a target tissue into distinct zones of gene expression, as originally proposed by the “French flag” model (Rogers & Schier, 2011; Wolpert, 1969). The genes triggered within each zone respond to the level of the activating transcription factor, such that different sets of genes are induced, accordingly (Sagner & Briscoe, 2019). While the morphogen levels decay in a continuous and graded manner over the cell field, distinct zones of gene expression marked by sharp borders are generated by the non-linear response of target genes to the morphogen. The capacity of cells to read small changes in morphogen levels is translated to an “all or none” transcriptional output, by virtue of cooperative responses to morphogen signaling.

While the morphogen dictates the division of a naive field into distinct uniform zones, there may be instances where additional patterning within these domains is necessary. For example, in response to the morphogen, coordinated cell movement and tissue morphogenesis may require manifestation of the graded positional information. This output is inherently distinct, in that cells within a given zone would respond differently according to the level of the morphogen. To examine if a morphogen gradient can also elicit patterning *within* a designated zone, we have turned to zygotic patterning along the dorso-ventral (DV) axis of the *Drosophila* embryo, where both aspects are operating.

Polarity along the DV axis of the *Drosophila* embryo is initially dictated by the Spaetzle/Toll pathway (Moussian & Roth, 2005). All the components of the pathway are contributed maternally. The ligand morphogen Spaetzle (Spz) is distributed in a graded manner within the extracellular peri-vitelline fluid which surrounds the embryo, with peak levels at the ventral midline (Haskel-Ittah et al., 2012; Rahimi et al., 2019). Following binding by the Spz ligand, activation of the Toll receptor leads to nuclear targeting from the embryo cytoplasm of the NFkB-related transcription factor, Dorsal, from the embryo cytoplasm, a process which is graded along the DV axis (Roth et al., 1989; Rushlow et al., 1989; Steward, 1989).

The *Drosophila* embryo is a syncytium during the initial 13 synchronous nuclear division cycles that follow fertilization. While Toll pathway activity and graded Dorsal nuclear accumulation are observed from nuclear cycle (NC) 11 onwards, with Dorsal re-entering the nuclei after the completion of each syncytial nuclear cycle. The critical manifestation of zygotic DV axis target-gene expression takes place, however, during the initial 45 minutes of the 14^th^ and final nuclear cycle, in parallel to embryo cellularization, during which all ∼6000 cortical nuclei are enclosed in individual cells. Due to nuclear envelope breakdown, Dorsal re-enters the nuclei after the completion of each syncytial nuclear cycle. Live measurements have shown that at NC 14, the graded Dorsal nuclear localization is re-established within 10-15 minutes (Rahimi et al., 2019).

The extracellular Spz gradient is thus translated to a nuclear-localization gradient of Dorsal, that dictates the onset of zygotic gene expression and the definition of the three DV axis zones within the blastoderm embryo: The ventral domain, expressing *twist* (*twi*) and *snail* (*sna*) will invaginate and give rise to the mesoderm. The lateral zone, expressing *rhomboid* (*rho*) and *short gastrulation* (*sog*) will comprise the neuroectoderm. Finally, the dorsal domain expressing *dpp* will generate the dorsal ectoderm and amnioserosa (Liberman et al., 2009; Moussian & Roth, 2005). Within each of these domains, the expression of the cardinal target genes appears uniform across the region.

In addition to defining broad zones along the entire axis, the same pathway operates within a single domain, the mesoderm, to elicit the coordinated three-dimensional cell shape changes and actomyosin-driven movement of mesodermal cells undergoing gastrulation (Heer & Martin, 2017; Martin et al., 2009). During gastrulation, the ventral cells expressing *twi* and *sna* invaginate by actomyosin-mediated constriction of their apical surfaces to form the ventral furrow. This constriction is driven by activation of Rho1, that leads to recruitment of the actin nucleator Diaphanous (Dia), as well as Rho kinase (Rock) which activates myosin II (Dawes-Hoang et al., 2005; Miao & Blankenship, 2020; Nikolaidou & Barrett, 2004; Perez-Vale & Peifer, 2020). Invagination of the mesoderm and ventral furrow (VF) formation, which involve the 1,500-odd ventral cells, take place in a coordinated fashion immediately after completion of cellularization. VF formation initiates as a wedge-shaped internalization of the ventral-most cells, a pattern that results from the graded recruitment of myosin (Denk-Lobing et al., 2020; Heer & Martin, 2017). In mutant backgrounds where myosin is recruited uniformly, the entire cell cohort invaginates simultaneously. Thus, it is necessary to control gene expression within the ventral-most region, in a manner that will lead to graded myosin recruitment.

Most of the proteins comprising the contractile machinery are provided maternally, and are therefore ubiquitous and uniformly distributed within the early embryo. The ability of the Toll pathway to shape the spatial pattern of mesoderm internalization, relies therefore on directing the patterned expression of zygotic target genes that will drive actomyosin contractility. Three key zygotic target genes that are expressed in the ventral part of the embryo, *folded gastrulation* (*fog*), *mist* and T48, are involved in myosin recruitment. They trigger the key initiating event, activation of Rho1, via its activator RhoGEF2 (Mason et al., 2016). Fog is a secreted ligand that activates G-protein coupled receptors (GPCRs), and Mist is the corresponding GPCR. Binding of Fog to Mist in ventral cells results in the activation and release of the membrane-tethered G*α* protein Concertina, leading, in turn, to recruitment from the cytoplasm and activation of RhoGEF2 (Manning et al., 2013; Perez-Vale & Peifer, 2020). In addition, the transmembrane protein T48, which is localized on the apical membrane, is also involved in RhoGEF2 recruitment to this cellular domain (Kolsch et al., 2007), where myosin II is subsequently required for apical cell constriction (Figure 1).

**Figure 1.**
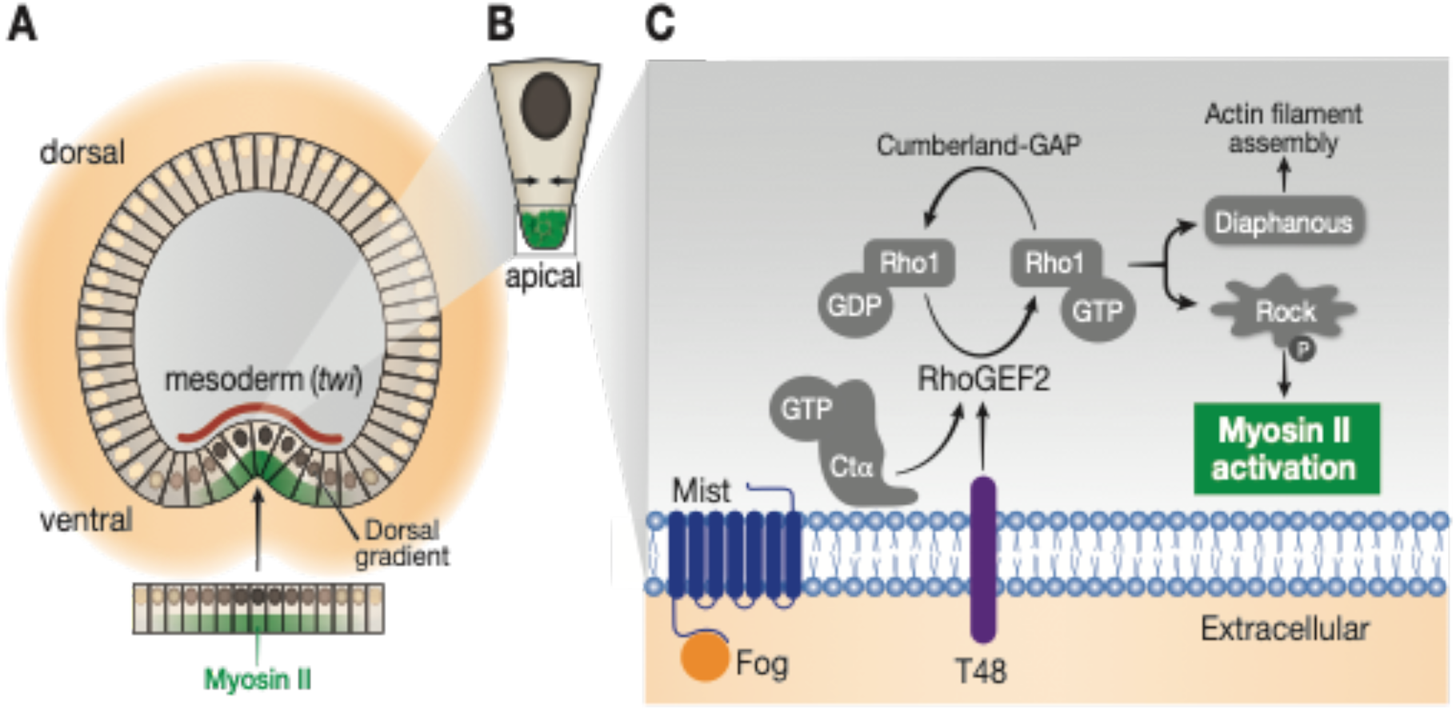
Graded activation of RhoGEF2 drives gastrulation. A) Schematic cross-section of an embryo initiating ventral furrow (VF) invagination during NC 14, after completion of cellularization. The nuclear gradient of Dorsal protein drives expression of *twist* in the mesoderm. Recruitment of Myosin II (green shading) to the apical surface of the ventral cells drives their apical constriction and invagination. B) The apical wedge-shape of the gastrulating tissue is triggered by graded recruitment of Myosin II, peaking at the ventral midline. C) Molecular basis of VF formation: Myosin II is recruited and activated apically by Rho1-GTP/Rock, following Rho1 activation by RhoGEF2. Two mechanisms combine to ensure the apical activity of RhoGEF2: Extracellular Fog ligand triggers the Mist G-protein coupled receptor to release the G*α* protein Concertina (Cta), while T48 is an apically-localized transmembrane protein that facilitates RhoGEF2 recruitment. All components marked in gray and Myosin II are deposited maternally. Expression of *fog, mist* and *T48* is induced zygotically, raising the possibility that their expression pattern is key to the graded recruitment of Myosin II.

In this work we analyzed the expression patterns of the zygotic genes *T48* and *mist*, encoding transmembrane proteins that regulate actomyosin contractility. We found that graded nuclear localization of Dorsal leads to a corresponding graded accumulation of mRNA for both genes through two complementary mechanisms. Activation of gene expression expands over time from the ventral region with high nuclear Dorsal to lateral regions where the levels are lower. Thus, nuclear Dorsal levels dictate the duration of gene activation. In addition, once a transcription site is primed, the rate of RNA Polymerase II (Pol II) loading depends on the level of nuclear Dorsal. A combination of these two responses contributes to the graded accumulation of *T48* and *mist* mRNA in the cytoplasm. In addition to the expression pattern of these two regulators, the timing of their activity is also critical, since the cell movements of gastrulation can commence only after the completion of cellularization. While formation of the mRNA gradients ensues during the initial stages of NC 14, the manifestation of their activity should nevertheless be delayed until cellularization is completed. We find that such timing is dictated by long introns found in the *T48* and *fog* genes, which delay the appearance of mature mRNAs that can undergo translation.

## Results

### Activation of *twist* expression follows a switch-like mechanism

*twist* (*twi*), encloding a bHLH transcription factor, is one of the two cardinal zygotic genes that (along with *snail*) define the mesodermal zone downstream of the Dorsal gradient, and subsequently drive the expression of a plethora of tissue-specific genes. To monitor the expression of *twi* quantitatively, we used single-molecule fluorescent *in-situ* hybridization (smFISH), which allows to visualize both the active transcription sites (TSs) in the nucleus, and the accumulation of individual mRNA molecules in the cytoplasm. Since the analysis is carried out on fixed embryos, each embryo represents a single time point, and the combined analysis of such “snapshots” of multiple staged embryos allows reconstructing the dynamic expression profile.

We find that *twi* expression displays a switch-like behavior, manifesting a sharp on/off threshold in response to Dorsal-nuclear localization levels. Active *twi* TSs appear simultaneously throughout the entire mesoderm region. The spatial distribution of these sites within the mesoderm is uniform along the DV axis (Figure 2 A-C). Most of the mesodermal nuclei display two TSs, indicating that both alleles are transcriptionally active. The median fluorescence intensity *of twi* TSs along the DV axis is also uniform, indicating a similar rate of Pol II loading onto all *twi* gene loci throughout the mesodermal domain (Figure 2 D). The uniform activation of *twi* transcription is also corroborated by a recent study that followed live accumulation of Twi protein, and identified early and even appearance of Twi protein throughout the mesoderm (Dufourt et al., 2020). The rapid activation of *twi* expression by Dorsal could be facilitated by Twi protein binding to the *twi* regulatory region. Identification of binding of Twi protein to the regulatory region of its own gene (J. Zeitlinger, personal communication, April 2021) may represent such a “Feed forward” mechanism, to enhance robust *twi* expression in nuclei where the levels of Dorsal are above the designated threshold.

**Figure 2.**
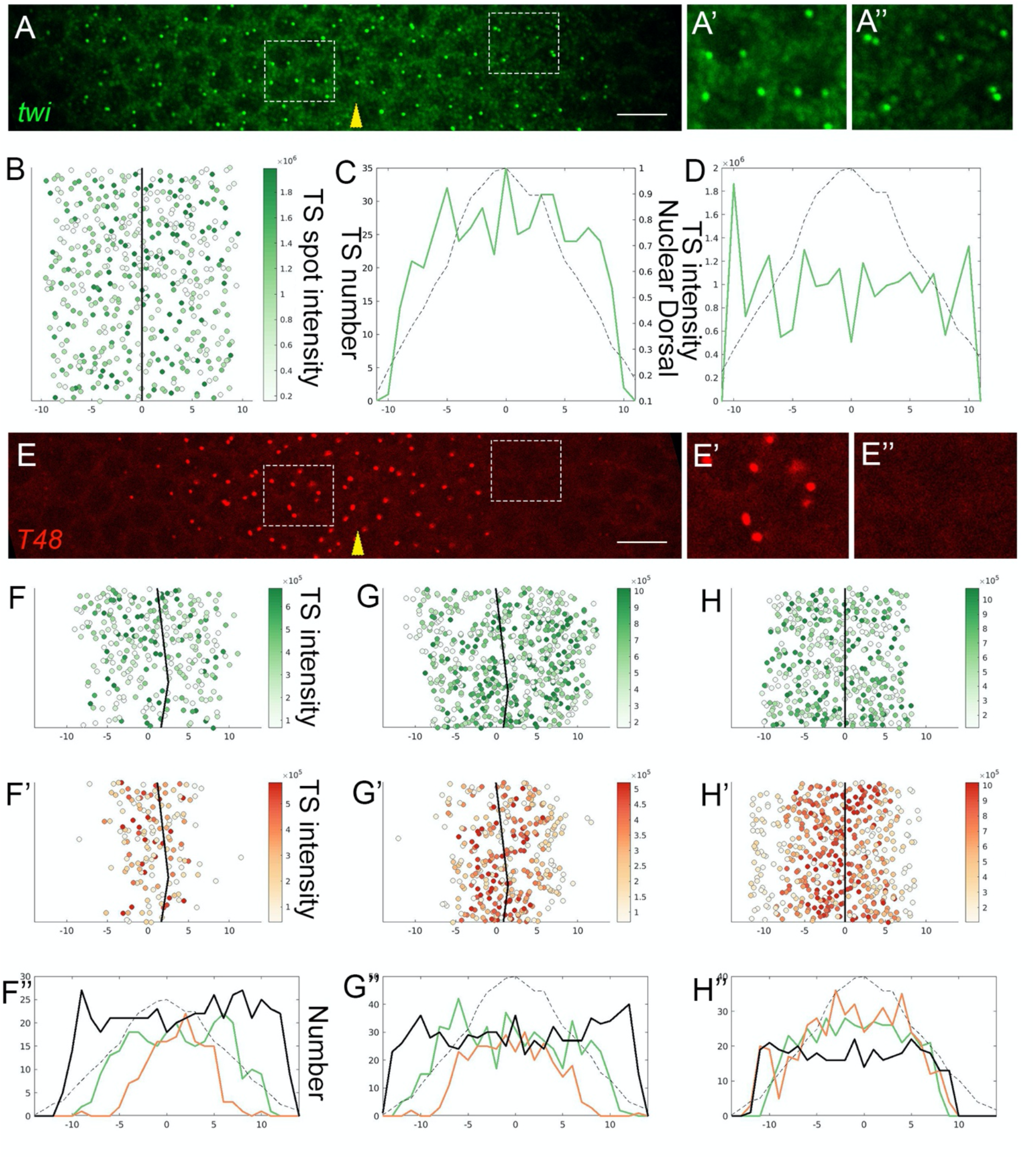
Dynamics of *twi* and *T48* transcription priming. A) *twi* expression in early NC 14 embryos was monitored by smFISH (green), and the transcription sites (TSs) were readily identified as prominent spots within the nuclei. Prominent *twi* expression was detected in all 20 nuclear columns comprising the future mesoderm. Most nuclei exhibited two TSs per nucleus. The ventral midline is marked by a yellow arrowhead. Scale bar 10 μm. Magnified views of a ventral and more lateral area within the mesoderm, marked by squares, are shown in panels A’ and A”, respectively. B) Quantitative measurement of the intensity of single *twi* TSs within the mesoderm of the embryo visualized in panel A. In this and all subsequent figures, 0 on the X axis marks the ventral midline, and numbers correspond to nuclear columns. The solid black line represents the ventral midline. n (TS number)=480. C) The number of *twi* TSs per column along the mesoderm is similar. In this and all subsequent figures, the dashed line marks the relative measured amount of nuclear Dorsal according to (Ambrosi et al., 2014). D) The relative median intensity of *twi* TSs is similar along the entire mesoderm domain. These results indicate that *twi* expression is activated by a switch mechanism, at a uniform level throughout the future mesoderm. E) Expression of *T48* (red) was monitored by smFISH in the same embryo shown in panel A. Scale bar 10 μm. Magnified views shown in panels E’ and E” correspond to the same regions shown in panels A’ and A”. While both *twi* and *T48* TSs are observed in the ventral domain, the lateral mesoderm was devoid of *T48* TSs. These results indicate that Twi is not sufficient for inducing *T48* expression. F-H”) Monitoring embryos of different ages (as defined by the number of *T48* TSs) within early NC 14, revealed the dynamic expansion of *T48* priming, covering eventually the entire *twi* domain. Shown are quantifications of *twi* TS intensities (F-H) and T48 TS intensities (F’-H’) for three embryos. Panels F”-H” compare the number of TSs per column of *twi* (solid green line) and *T48* (solid orange line) in these embryos. The solid black line denotes total nuclei number, monitored by DAPI staining. F-n=283. F’-n=155, G- n=565, G’ n=321, H-n=414, H’-n=471.

### Triggering graded *T48* expression

To examine the basis for graded actomyosin contractility within the mesoderm, we followed the expression of key zygotic target genes. Towards this end, we chose to focus on T48 and Mist, given the central role which these transmembrane proteins play in eliciting an actomyosin response to the DV patterning program.

We followed the expression of *T48* and *twi* in parallel, in the same embryos. While *twi* expression encompasses the entire mesoderm from the outset, the expression of *T48* was initially observed only in the central (ventral-most) part of the mesoderm (Figure 2 E). In older embryos, however, the distribution of TSs for *twi* and *T48* showed a complete overlap (Figure 2 F-H). We conclude that while *twi* expression is triggered simultaneously in all mesodermal nuclei, the onset of *T48* expression expands over time. Temporal expansion of *T48* expression has also been observed by live imaging using MS2 tagging (Lim et al., 2017). Twi protein is necessary for activation of *T48* expression, since no transcription of *T48* takes place in *twi*-mutant embryos (Leptin, 1991). The distinct temporal profiles of *twi* and *T48* expression imply, however, that additional factors besides Twi regulate *T48* expression. The most likely candidate for such a factor is Dorsal, whose nuclear levels within the mesoderm are graded. This nuclear distribution represents the final asymmetry that is induced by the Toll pathway along the DV axis. Binding sites for both Twi and Dorsal have been identified on the *T48* enhancer (J. Zeitlinger, personal communication, April 2021).

From our analysis of the number of *T48* TSs that are activated in each nucleus at different time points, we concluded that the graded activation reflects the independent response of individual *T48* loci, since many nuclei display only one active TS. Furthermore, ventral nuclei exhibited primarily two active *T48* TSs per nucleus, whereas more lateral nuclei displayed only one TS. In other words, depending upon the level of Dorsal, there is a certain independent probability of “priming” each *T48* locus, so as to make it permanently accessible for the recruitment of transcription factors and RNA polymerase II. Over time, nuclei start to display two active TSs, first in the ventral region and subsequently in more lateral domains. This pattern is distinct from that of *twi* TSs, which were mostly represented as two per nucleus throughout the future mesoderm (Figure 3 A-C).

**Figure 3.**
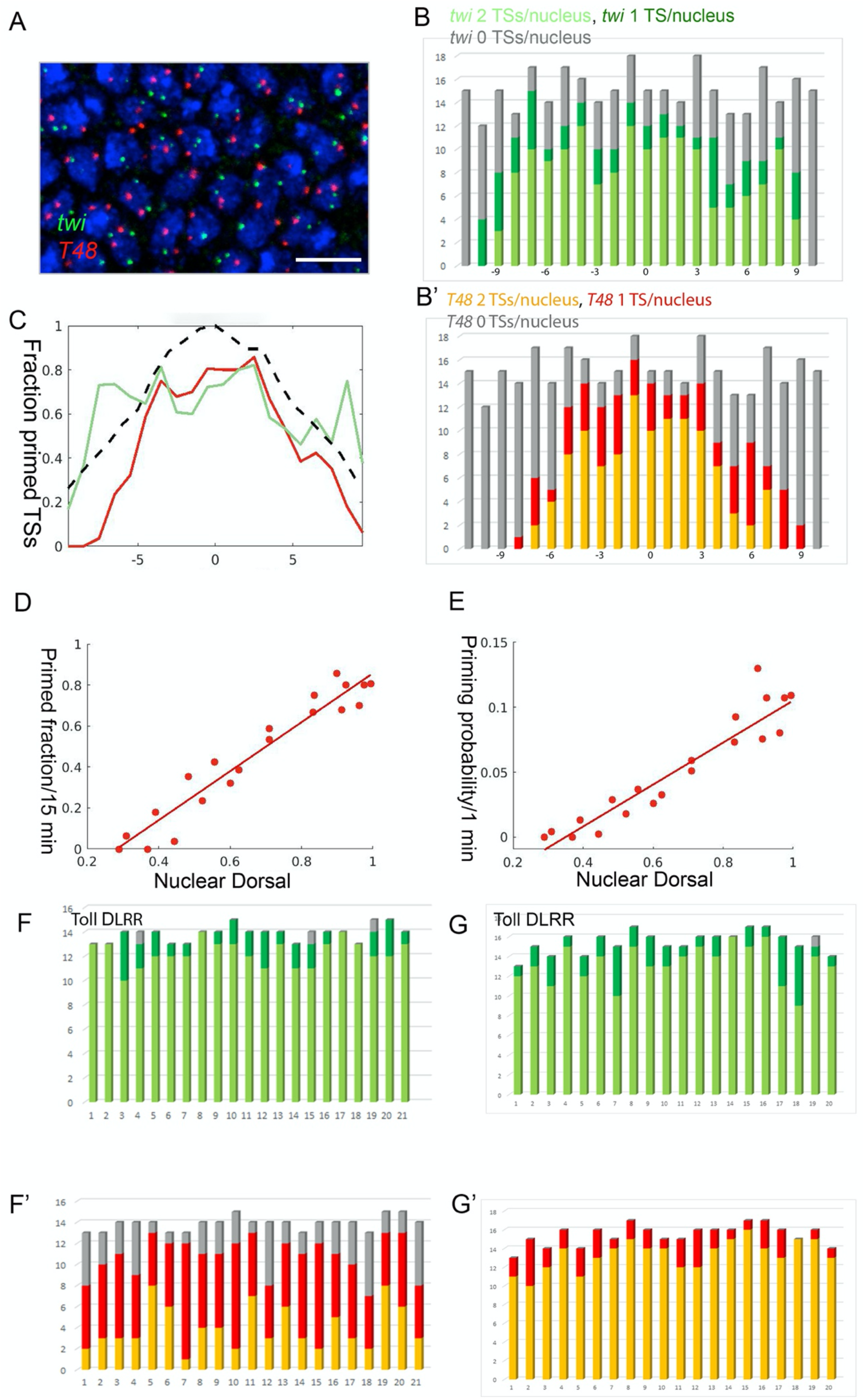
The probability of *T48* TS priming depends on Dorsal levels. A-C) Quantification and graphical analysis of TSs in an early NC 14 embryo probed for *twi* (green) and *T48* (red). Nuclei (panel A) visualized with DAPI (blue). The probability of TS activation was gauged from the number of TSs activated in each nucleus. While most nuclei within the mesoderm exhibited 2 *twi* TSs per nucleus (e.g. fully primed), a gradient of *T48* activation was apparent. Most of the ventral nuclei displayed 2 TSs, while the lateral nuclei showed 1 or no TSs. Identification of many nuclei with only one *T48* TS indicates that the probability of priming transcription operates independently for each of the two loci within one nucleus. Scale bar 10 μm. D) The primed fraction of loci is linearly correlated to the nuclear Dorsal level (correlation 0.96). E) The embryo was estimated to be exposed to a stable Dorsal gradient for ∼15 minutes within NC 14. This time window can be used to calculate the probability of TS priming along the future mesoderm, which is again linearly correlated with the level of nuclear Dorsal (correlation 0.94). F,G) Uniform expression of the constitutively active Toll^ΔLRR^ construct drives activation of the pathway and nuclear targeting of Dorsal along the entire embryo circumference. Induction of *twi* and *T48* TSs responds accordingly. In a younger embryo (F), two *twi* TSs are observed in most nuclei, while only a single *T48* TS is observed in a large fraction of nuclei. In an older embryo (G) the majority of nuclei exhibit two TSs for both genes.

The embryo in Figure 3 A-C is ∼ 25 minutes into cycle 14, as estimated from the size and shape of the nuclei (Lecuit et al., 2002). Since establishing a graded pattern of nuclear Dorsal requires 10-15 minutes following the onset of NC 14 (Rahimi et al., 2019), the nuclei of this embryo were exposed to the Dorsal gradient for ∼15 minutes. The fraction of nuclei displaying activated *T48* TSs was measured along the ventral domain of this embryo. This value can then be used to derive the priming probability per minute at different positions along the DV axis (Figure 3 D,E).

Using this approach, we find that the probability of *T48* TS activation peaks at the ventral midline, and declines towards the lateral region. The probability distribution resembles the shape of the Dorsal-nuclear gradient. This tight correlation (Pearson’s Rho (p):0.94) indicates that the probability of TS activation declines linearly with the reduction in Dorsal levels. Such a monotonic response to Dorsal is in sharp contrast to the non-linear response of the *twi* enhancer to Dorsal, which generates the sharp borders of *twi* transcription.

The instructive role of nuclear Dorsal in determining the probability to trigger the *T48* loci was examined in embryos where the Toll pathway was ubiquitously activated by uniform expression of the constitutively active Toll^ΔLRR^ construct (Rahimi et al., 2016). Nuclei around the entire embryo circumference exhibited a similar response. In younger embryos, two *twi* TSs per nucleus appear in most nuclei, while mostly one or (in a minority) two *T48* TSs were observed. This intermediate response of *T48* TSs reflects the probability of priming, which requires some time to result in a significant fraction of active TSs. Indeed, in older Toll^ΔLRR^ embryos most nuclei presented two active TSs for both *twi* and *T48* (Figure 3 F,G).

### Graded recruitment of Pol II to the *T48* gene

Median intensity of the individual *twi* TSs across the mesoderm appears constant (Figure 2 D), indicating a similar time of onset and a comparable rate of Pol II loading. Measuring and analyzing the intensity of *T48* TSs as a proxy for the number of polymerases traversing the gene is more complex, given the gradual, time-dependent progression of expression initiation of this gene along the DV axis. It is imperative, therefore, to only consider those *T48* loci that are already at a steady-state, and are producing mature transcripts. We therefore utilized distinct smFISH probes for the 5’ and 3’ domains of *T48*, and limited our analysis of the intensity of *T48* TSs to loci that displayed both a 5’ and a 3’ probe signal, indicating that some Pol II molecules had reached the 3’ region of the gene (Figure 4 A-D). Monitoring the intensity of the 5’ probe for these sites, provided a cumulative measure for the number of polymerases along the gene. A graded median intensity gradient was observed across the ventral region (Figure 4 C). A tight correlation of the Pol II loading rate with the position (and hence on nuclear Dorsal levels) was also detected (p=0.47) (Figure 4E).

**Figure 4.**
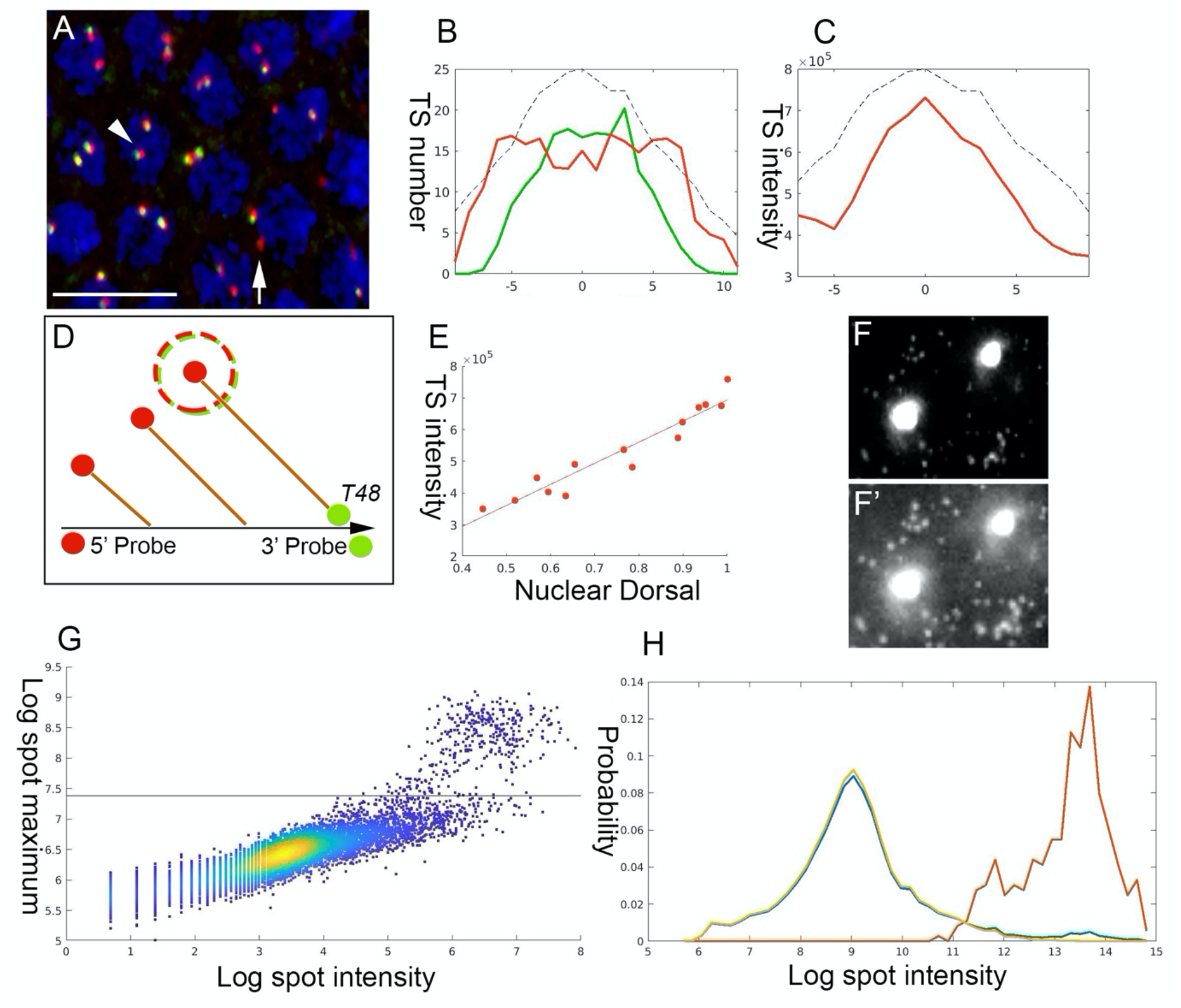
The rate of Pol II loading on *T48* depends on Dorsal levels. A) A region within the mesodermal domain of an early NC 14 embryo hybridized to two distinct *T48* smFISH probes directed against either the *T48* 5’ region (red) or the *T48* 3’ domain (green). TSs with initiated transcription that has not yet reached the 3’ end appear as red dots (arrow), while TSs that are producing mature transcripts are yellow (arrowhead). Nuclei are visualized with DAPI (blue). Scale bar-10 μm. B) Quantification of TS numbers in this embryo reveals that T48 transcription has been initiated to a similar extent throughout the mesoderm (red), Pol II has not yet reached the 3’ end of the gene in a significant number of TSs in the lateral region (green). C,D) The *T48* Pol II loading capacity along the mesoderm was followed by monitoring the median intensity of the 5’ probe (red), only in those TSs that also exhibit a signal for the 3’ probe (marked as a dashed circle in D). E) A distinct dependence of the loading rate on the position along the DV axis (and hence Dorsal level) was observed (p=0.47). F,F’) *T48* signal intensity was followed with a smFISH probe covering the entire coding sequence (gray). The prominent spots in the nuclei represent the TSs, while the weak spots in the cytoplasm correspond to individual mRNA molecules. Different levels of exposure of the same image highlight the two types of signals. G) The maximal pixel (y-axis) and the cumulative pixel intensity (x-axis) of each spot were compared. The rationale is that single or cytoplasmic clusters of mRNA molecules would be present in large numbers and display a lower maximal intensity, due to their lower concentration. The TSs would be marked by high maximal intensity due to the elevated local concentration of mRNA molecules. The horizontal black line distinguishes between the two populations. H) Cumulative spot intensity histogram identifies two peaks, corresponding to the median single mRNA intensity and TS intensity, respectively (total number-blue, mRNA-yellow, TS-red). The median intensity of the TSs was 70 times higher than that of the single mRNA, providing an estimate of the average *T48* Pol II loading. Taking the size of the *T48* gene into account, this value is ∼ten-fold lower than the maximal Pol II loading capacity, indicating that differential Pol II loading is likely within the linear range.

To determine the average number of Pol II complexes that are loaded on *T48*, we compared the intensity of nuclear TSs with that of individual mRNA spots in the cytoplasm, using a probe covering the entire coding sequence (Figure 4 F). The cumulative and maximal pixel intensity and volume of each spot were monitored. The rationale for this approach is that the cumulative spot intensity indicates the number of mRNAs, while the maximal intensity distinguishes highly concentrated mRNA molecules at the TS in the nuclei, from the more dispersed mRNA clusters in the cytoplasm. A histogram across the spot intensity (in Log scale) for each of these groups identified two peaks, corresponding to the median single mRNA intensity and TS intensity, respectively (Figure 4 G,H). The median intensity of the TSs was 70 times higher than that of the single mRNAs, providing an estimate for the average extent of Pol II loading onto *T48* loci (Figure 4 G,H). Since the *T48* gene is 29 kb long, this translates to a density of one Pol II complex every ∼400 bp, which is about ten-fold lower than the maximal loading that is physically possible. Thus, control of the number of Pol II molecules loaded on the *T48* locus is likely within the linear dynamic range.

### Dual mechanism leading to a graded *T48* mRNA accumulation

A combination of the two mechanisms outlined above, graded onset of TS priming, and graded polymerase loading rates of the gene across the ventral region, should give rise to a graded distribution of *T48* mRNA. Indeed, quantitation of *T48* mRNA in the cytoplasm showed a graded pattern that is highly correlated with nuclear Dorsal levels (p=0.92+-0.02) (Figure 5 A-C). Since Twi distribution in the mesoderm is uniform, we assume that nuclear Dorsal underlies the graded expression of *T48*.

**Figure 5.**
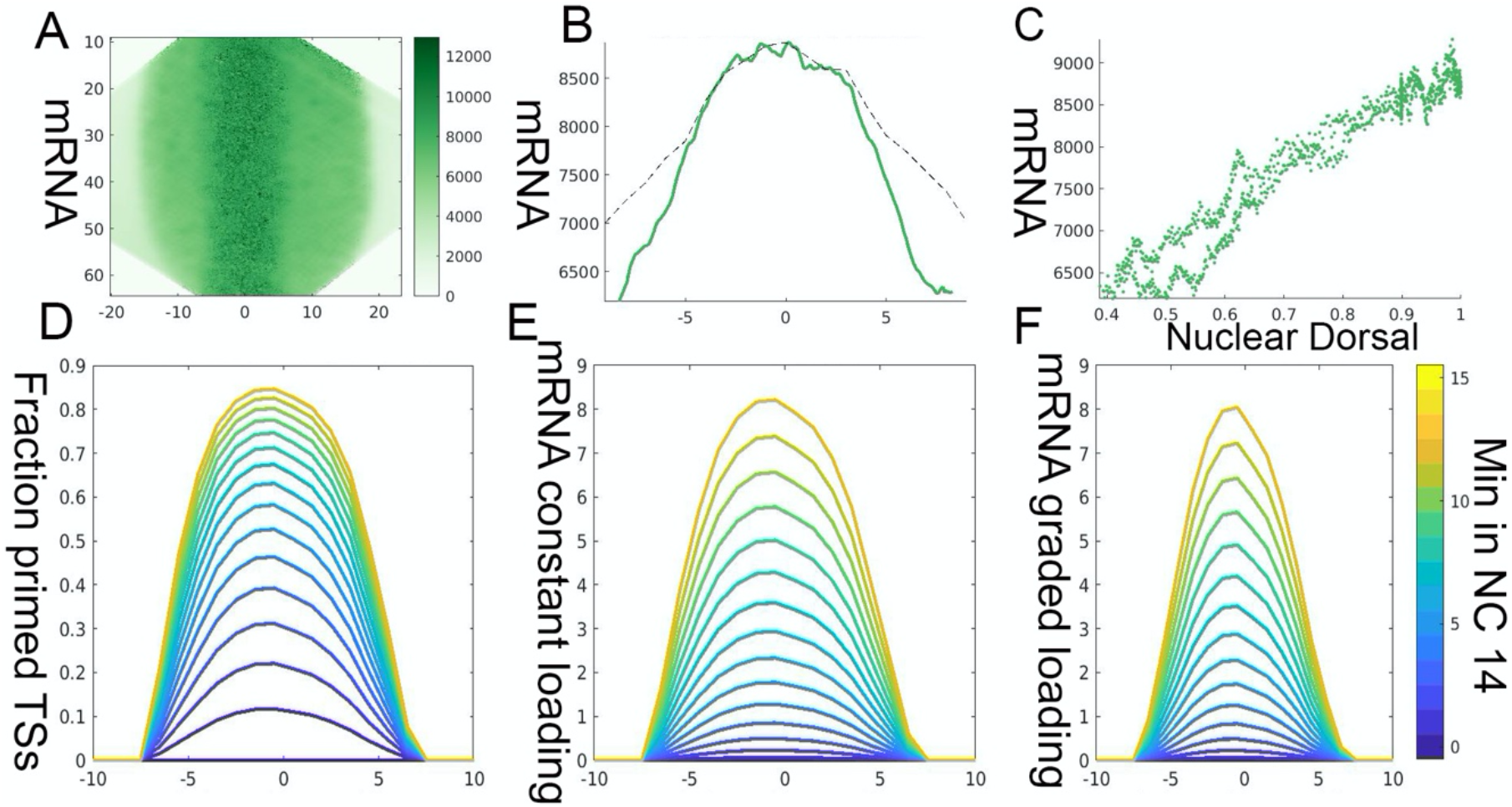
Priming probability and graded Pol II loading generate graded *T48* mRNA accumulation. A,B) To follow the pattern of *T48* mRNA cytoplasmic accumulation and omit TS spots, the median pixel intensity at each distance from the midline was calculated for an embryo in which *T48* TSs could be observed at the lateral edges of the mesoderm. A graded accumulation of *T48* mRNA along the future mesoderm is readily apparent and is correlated with the levels of nuclear Dorsal. C) The level of cytoplasmic *T48* mRNA correlates with the distance from the ventral midline and nuclear Dorsal (p=0.95). D) Based on the probability of *T48* TS priming that was calculated from experimental data (see Figure 3), the fraction of primed nuclei over time was simulated. E) If we assume that the rate of Pol II loading onto primed gene loci is similar throughout the mesoderm, a gradient of mRNA accumulation will still be generated, since ventral loci are primed earlier than more lateral ones. F) The resulting mRNA gradient sharpens further when we include a graded Pol II loading rate in the simulation. The combined effect of Dorsal-dependent priming probability and graded loading, gives rise to graded accumulation of *T48* transcripts in the cytoplasm.

The quantitative measurement of *T48* TS priming (Figure 3 D,E) allows to simulate the dynamic pattern of production and accumulation of mRNA (Figure S1). The fraction of primed TSs increases over time in a graded manner. If every primed site is transcribing at a similar rate, this would still lead to graded accumulation of mRNA, since the ventral TSs were primed earlier (Figure 5 D,E). We assume that the transcribed mRNAs are stable within the relevant time window.

In addition to the temporal order of TS priming, the actual rate of Pol II loading depends on the level of nuclear Dorsal. When this aspect is added to the simulation, the shape of the accumulating mRNA peak becomes sharper (Figure 5 F). Since the priming of TSs reaches saturation at the midline, i.e. almost all TSs become primed, this additional feature contributes to the shape of the mRNA peak within the relevant time window. It is interesting to note that the probability of TS activation represents a compromise between two opposing considerations. Saturation would not be reached at low priming probabilities, but the actual levels of mRNAs that are produced may be too low if only a small fraction of TSs would be activated. It appears that the system is utilizing an optimal probability range, to produce within the relevant time window sufficient, yet graded, mRNA levels.

### Graded accumulation of *mist* transcripts

Expression of *mist* was analyzed in a similar way to *T48*, and exhibited comparable features. Binding sites for both Twi and Dorsal have been identified on the *mist* enhancer (J. Zeitlinger, personal communication, April 2021). Graded priming of TSs was identified over time, where the initial activation took place along the ventral midline (Figure 6 A,B). Monitoring TS intensity revealed a gradient of *mist* TS intensities, reflecting a position-dependent Pol II loading rate (Figure 6 A’,B’). Of note, as *mist* is a relatively short gene (9.3 kb), we assumed transcription to reach steady state on all TSs in late embryos. *mist* mRNA accumulated in the cytoplasm in a graded manner (Figure 6 A’’,B’’). Monitoring the number of activated TSs per nucleus revealed more nuclei with two TSs in the ventral region (Figure 6 C). Finally, uniform activation of the pathway following expression of Toll^ΔLRR^ resulted in a uniform level of *mist* priming along the DV axis (Figure S2).

**Figure 6.**
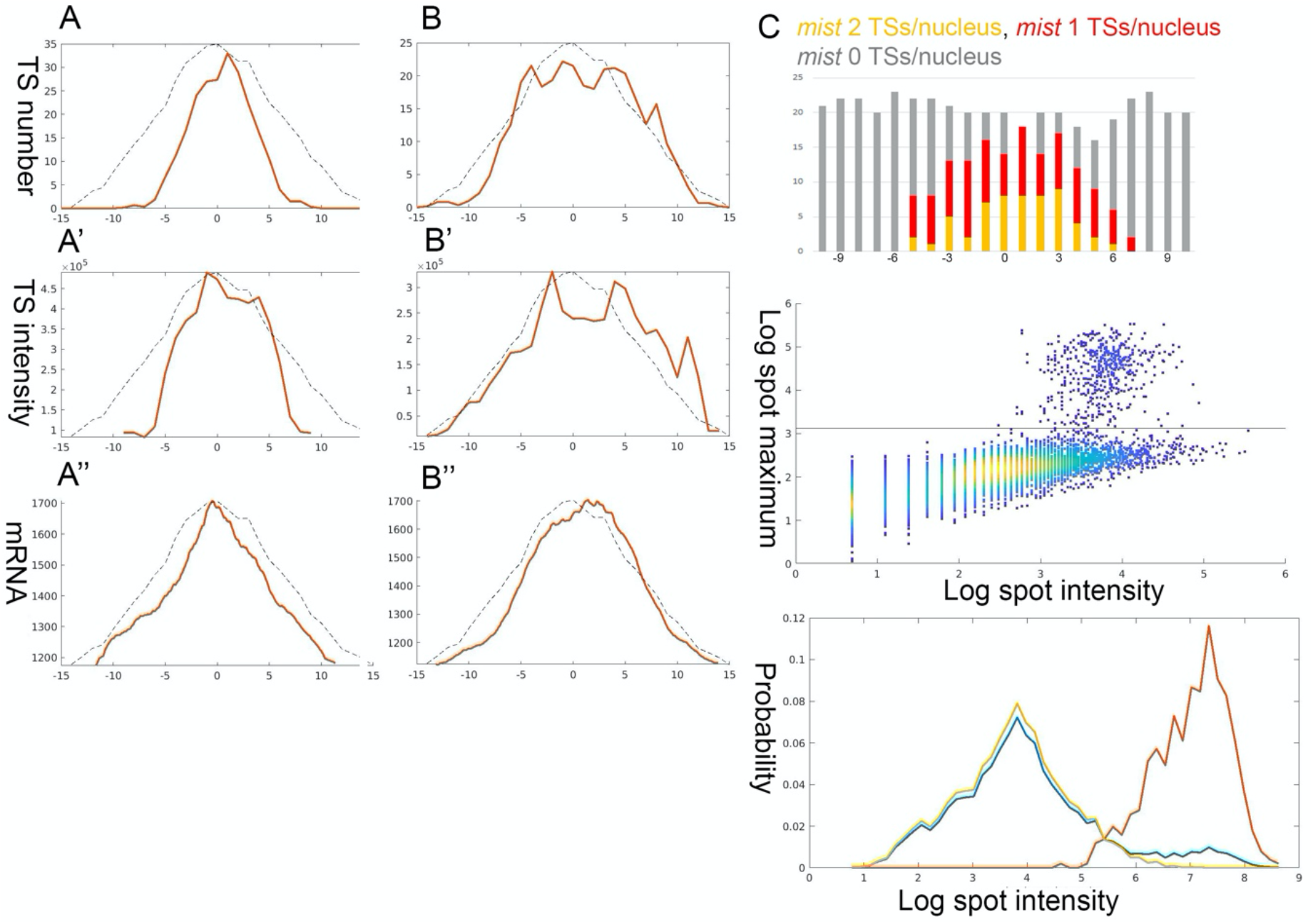
Graded accumulation of *mist* transcripts. Mist, a GPCR, is the second zygotically-expressed transmembrane protein that participates in RhoGEF2 activation and recruitment. Quantification and graphical analysis of *mist* expression during early NC 14 was performed using a single 5’ region smFISH probe. A-B”) Expression of *mist* in a pair of embryos displays similar hallmarks to those identified for *T48*. Dynamic *mist* priming was seen by the lateral expansion of TSs over time (the embryo analyzed in panels A-A” is younger than the one analyzed in panels B-B’). Pol II loading depends on Dorsal levels, as indicated by the graded TS intensity (p=0.21). The resulting mRNA accumulation of *mist* displays a sharp graded pattern (p=0.95). C) The Dorsal-dependent priming probability of *mist* can be assessed by monitoring the number of activated loci per nucleus. D,E) The average number of Pol II complexes per *mist* locus was calculated as 30. Taking the gene size into account, this corresponds to an average density that is 1.4 times higher than the one calculated for *T48*, and is still well within the linear range.

To determine the Pol II loading rate onto the *mist* gene loci, the intensity of TSs and single mRNA spots was compared (Figure 6 D,E). From this, the median number of Pol II complexes per gene is estimated at 30. Taking the size of the gene into account (9.3 kb), this distribution is 1.4 times denser than the one calculated for *T48*, but still well below saturation levels.

### Intron delay of *T48* expression

The graded accumulation of mRNAs for the *T48* and *mist* genes is effectively driven early in NC 14, following completion of the final syncytial cycle of nuclear divisions. However, the protein products that would trigger apical actomyosin contractility, including RhoGEF2 (Mason et al., 2016) and Myosin II (Dawes-Hoang et al., 2005), are recruited apically and function only after cellularization is complete, 30-40 minutes after the onset of NC 14 (see also Movie S1), necessitating a delay. An indication that regulation of mature mRNA production may contribute to this timing mechanism was observed when comparing the cytoplasmic mRNA accumulation patterns of *twi* and *T48*. As shown above (Figure 2), early NC14 embryos display both *twi* and *T48* active prominent TSs. However, while *twi* mRNA was readily detected in the cytoplasm of such embryos, no *T48* mRNA was observed (Figure 7 A). Since *twi* is a short gene of 2.2 kb, no delay in completion of transcription is expected to take place. In contrast, the *T48* and *fog* genes are ∼30 kb long, due to the presence of large introns, encompassing ∼25 kb. This organization is exceptional for early zygotic genes, which are typically intron-less or have only short introns (Artieri & Fraser, 2014). Given a transcription rate of ∼2 kb/min (Fukaya et al., 2017), such long introns could provide a delay that would allow the production of mature mRNA and protein only after the completion of cellularization.

**Figure 7.**
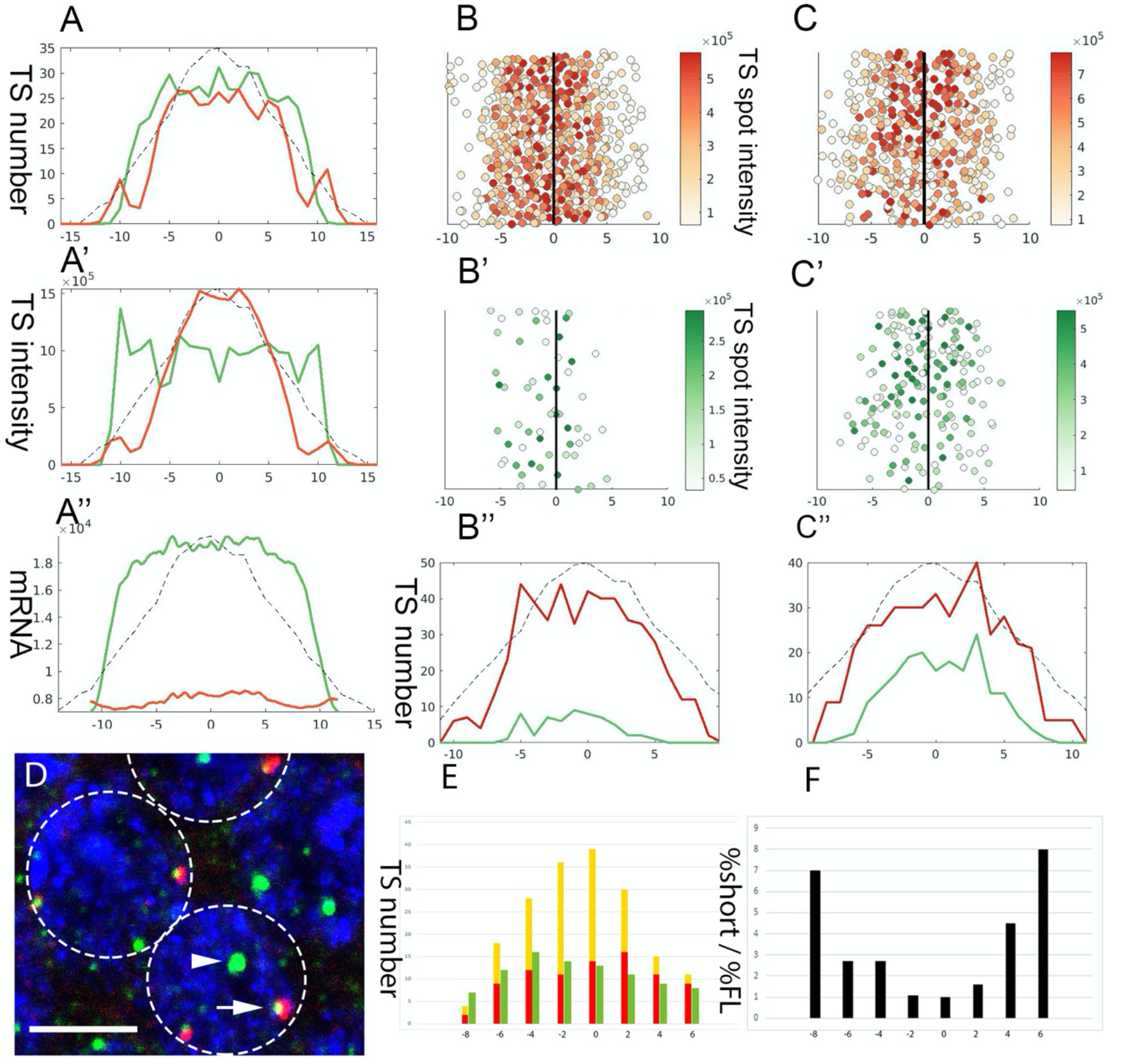
Intron delay dictates the timing of *T48* translation. While *T48* transcription may generate a graded distribution of mRNA shortly after the onset of NC 14, the actual recruitment of myosin II and apical constriction takes place only after completion of cellularization. What mechanism accounts for this timing? A-A’’) Expression of *twi* (green) and *T48* (red) was monitored simultaneously in the embryo analyzed in Fig. 2. *twi* mRNA accumulation was prominent. While not all nuclei exhibit *T48* TSs at this stage, the TSs at the ventral region were prominent. Yet, no *T48* mRNA was detected. We hypothesized that the unusually large intron of *T48* (20 kb) may account for the delay in transcript maturation. B,C) Probing embryos with a *T48* 5’ probe (red) and 3’ probe (green) distinguishes between TSs that have initiated transcription vs. ones that are already producing mature transcripts. In a relatively young embryo, as assessed by the total number of *T48* TSs (B-B”), the 5’ probe was detected in the entire ventral domains, while only sparse spots were monitored by the 3’ probe, primarily in the ventral-most region. The majority of Pol II complexes within the active *T48* loci have therefore not reached their 3’ end. In an older embryos (C-C’’) the 5’ and 3’ signals were overlapping. This difference underscores the delay in *T48* transcript maturation, that may stem from the time required to transcribe the long intron. B-n=509, B’-n=58, C-n=426, C’-n=184. D) To directly asses the contribution of the intron, a *T48* cDNA transgene that is driven by the *T48* enhancer was generated. Since the transgene is inserted at a different chromosomal position, it should display a TS signal distinct from the endogenous *T48* loci. To distinguish between the endogenous and cDNA transcripts, embryos were probed with a *T48* intron probe (red) and 3’ probe (green). We expect the endogenous TSs to be marked by both probes (arrow), while the cDNA should only react with the 3’ probe (arrowhead). Nuclei are marked by DAPI (blue). Scale bar 5 μm. E) The number of mature endogenous TSs (yellow) peaks at the ventral midline of an intermediate stage embryo, while the lateral region harbors transcripts that have reached the intron but not the 3’ end (red). In contrast, a fairly uniform distribution of cDNA TSs was observed (green), indicating that the cDNA TSs matured earlier than the endogenous loci. F) Plotting the ratio between the relative fraction of mature cDNA vs. endogenous TSs along the mesoderm demonstrates the faster maturation of the cDNA TSs.

### Movie S1

#### Dynamics of Myosin II recruitment

The dynamic localization pattern of Myosin II during early NC 14, monitored using a GFP “protein-trap” in the Myosin II heavy-chain (*zipper*) gene locus, as visualized in cross section by Lightsheet microscopy. During the process of cellularization, Myosin II is initially recruited to the basal aspect of the future cell membranes as they extend from the surface. Upon completion of cellularization, Myosin II disappears form the basal position and is recruited to the apical side in the ventral-most cells, where it drives furrow formation. The graded pattern of Myosin II recruitment within the ventral domain shapes the pattern of cell invagination. The movie was obtained at a rate of 1 frame/min.

If the intron prolongs *T48* transcription, we should expect spatial and temporal differences in the detection of TSs, using distinct *T48* 5’ and 3’ probes. Indeed, when embryos were simultaneously monitored with such probes, a clear delay was observed. In younger embryos the TSs (of variying intensity) reacting with the 5’ probe encompassed the entire mesoderm, while the 3’ probe marked only very few TSs, specifically in the ventral region (Figure 7 B). In older embryos most mesoderm nuclei displayed TSs reacting with both probes (Figure 7 C). This behavior demonstrates a substantial temporal delay between the onset of transcription and the time required for Pol II to reach the 3’ end of the locus and complete transcription.

If the delay is caused by the transcription of through the *T48* intron, then transcription of an intron-less *T48* gene within a given nucleus should be completed earlier than that of the endogenous locus. To test this prediction we fused a *T48* cDNA construct to its 5’ regulatory region (Lim et al., 2017), and generated transgenic flies which integrated the construct at a locus that is distinct from the endogenous one. By using distinct intron and 3’ probes, we could distinguish the completion of transcription at endogenous loci by their reactivity with both probes, whereas the cDNA locus reacted only with the 3’ probe (Figure 7 D). Analysis of the distribution of both TS types indeed demonstrated that completion of transcription for the cDNA TSs appears earlier than termination at the endogenous *T48* locus. While the mature endogenous TSs were detected mostly in the ventral domain, TSs exhibiting the cDNA 3’ domain only, were distributed roughly uniformly throughout the mesoderm (Figure 7 E). Accordingly, the fraction of TSs for the cDNA at the lateral domain was higher (Figure 7 F).

## Discussion

### Graded response to a morphogen gradient

The primary hallmark of morphogen gradients is the ability to utilize a graded morphogen distribution for the induction of distinct zones of gene expression. The ability to generate sharp borders of gene expression by reading small differences in morphogen levels, may necessitate cooperative binding mechanisms of the transcription factor(s) activated by morphogen signaling. We set out to explore if the information conveyed by the morphogen gradient can also be utilized to generate a graded response within the target tissue. In situations where the morphogen gradient is the only symmetry-breaking cue, morphogenetic movements of distinct zones in the target tissue may also require a graded transcriptional response to the morphogen.

We examined this paradigm during the establishment of the DV axis in the *Drosophila* embryo. The gradient of nuclear localization of the Dorsal protein, represents the only asymmetric cue that sets up the entire zygotic behavior of the embryo along this axis. An expected consequence of this gradient is the definition of three distinct domains (mesoderm, neuroectoderm and dorsal ectoderm) along the axis. In addition, the future mesoderm undergoes graded constriction by an actomyosin network, in order to fold and internalize in a wedge-shaped pattern. If this latter behavior is guided by a zygotic mechanism in the embryo, its origins may well originate in a mesoderm-specific transcriptional response to the Dorsal-nuclear gradient.

To examine the underlying transcriptional mechanism for graded actomyosin constriction, we focused on two zygotic genes encoding transmembrane proteins that regulate the activation of the small GTPase Rho1, a key initiating event in the process. T48 was shown to recruit RhoGEF2 to the apical surface (Kolsch et al., 2007), and Mist is a GPPCR that is activated apically by the ligand Fog, and releases the G*α* protein Concertina that activates RhoGEF2 (Manning et al., 2013; Mason et al., 2016; Perez-Vale & Peifer, 2020) (Figure 1). The *fog* gene is also zygotic, but since it encodes a protein that is secreted to the extracellular milieu of the peri-vitelline fluid, the timing of its expression, rather than the pattern, may be more relevant. All other elements in the actomyosin network are provided as maternal transcripts, and are ubiquitously expressed in the embryo.

We found two distinct Dorsal-dependent mechanisms that generate graded mRNA accumulation for *T48* and *mist*. Initially, expression of both genes is triggered only in the central (ventral-most) of the mesoderm, expanding laterally over time. This expansion is consistent with a TS priming response to the Dorsal-nuclear gradient, which is maximal at the ventral midline. Distribution of the number of activated TSs per nucleus indicates that the probability of priming is independent for each allele. Thus, two alleles within the same nucleus that experience similar levels of nuclear Dorsal, may be switched on at different times. The probability of TS priming is linearly correlated with the levels of nuclear Dorsal. This is in contrast to the sharp expression borders for the genes that define the three major zones along the embryonic DV axis, likely resulting from a cooperative response to Dorsal levels.

The molecular nature of the Dorsal-dependent mechanism that leads to priming of the *T48* and *mist* TSs is not known. One possibility is that alteration in the nucleosome organization could make the gene more susceptible to subsequent loading of Pol II complexes. Alternatively, since these enhancers also display binding sites for Twi, a stable association between Twi and Dorsal proteins may permanently alter the future accessibility of the site to incoming Pol II. Binding of Dorsal and Twi to target genes such as *sim*, was shown to facilitate their transcription, by enhancing the probability of DNA binding by the Notch intra-cellular domain (Falo-Sanjuan et al., 2019).

The different transcription onset times are expected to generate graded mRNA accumulation, with higher levels adjacent to the nuclei that were induced earlier. The effectiveness of this mechanism is restricted to a defined time window. When over time most of the transcription sites in the ventral region are triggered, the differences in mRNA production diminish. For optimal function, the probability of activation should be high enough to prime a significant number of TSs, and yet not plateau within the critical time window.

Once a TS is primed, the rate of Pol II loading is also dependent upon nuclear Dorsal levels. We can imagine a scenario where binding of Dorsal to the enhancers has on and off rates, such that the fraction of time that Dorsal is bound would depend on its concentration. The different Pol II loading rates would also contribute to the generation of an mRNA gradient. Furthermore, this mechanism will continue to operate even after the priming of TSs begins to plateau. Simulations based on our measurements, show that a combination of the two mechanisms can lead to a robust gradient of mRNA accumulation. Graded alteration in transcriptional bust size has also recently been demonstrated for BMP-target genes in the dorsal ectoderm of the *Drosophila* embryo (Hoppe et al., 2020). However, it is not clear if this graded response has a biological role. It is interesting to note that the utilization of two parallel regulatory mechanisms is also operating for patterning of actomyosin constriction. T48 guides RhoGEF2 apical recruitment, while Mist activation dictates the actual activation of RhoGEF2. We assume that this combination can make the resulting activation gradient of Rho1 sharper. The proposed mechanisms underlying graded activation of transcription by Dorsal are summarized in Figure 8.

**Figure 8.**
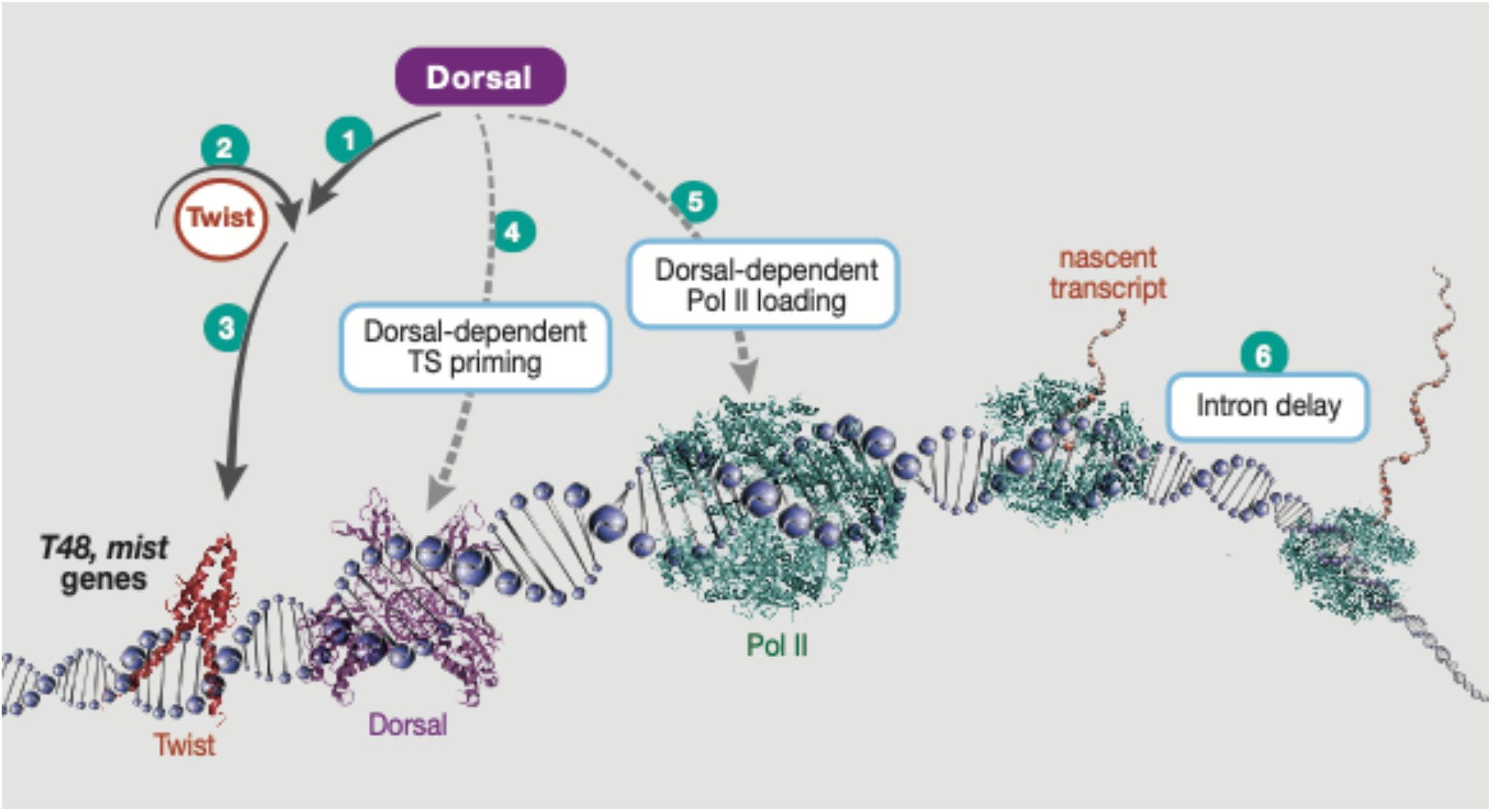
Graded accumulation of *T48* and *mist* transcripts drives gastrulation. A schematic view of the *T48* and *mist* genes, depicting proposed inputs from DV axis elements and gene transcription machinery, leading to graded mRNA accumulation in the mesoderm. The nuclear Dorsal protein gradient is the only source of asymmetry along the DV axis of the early *Drosophila* embryo. 1) Dorsal binding to the *twi* enhancer triggers expression with a sharp threshold. 2,3) The autoregulatory capacity of Twi assures robust and uniform *twi* expression along the future mesoderm. 4) Twi is also essential for *T48* and *mist* expression, but it is not sufficient. In order to prime expression from these loci, Dorsal is also required. Progressive induction over time depends on Dorsal nuclear levels. The probability of priming is dictated by the limiting levels of Dorsal, and will therefore progress over time from the ventral to the lateral region. 5) Once primed, loading of Pol II also depends on the Dorsal binding. Thus, the transient binding of Dorsal that is dictated by its nuclear concentration, will modulate the transcription burst size or frequency. A combination of both of these mechanisms will lead to a graded accumulation of mRNA. 6) Finally, generation of mature, translation-ready transcripts of the proteins leading to Myosin II recruitment can be influenced by the intron size of the genes. Unusually long introns in *T48* (and *fog*) provide a delay in transcript maturation, thereby providing a timing mechanism that ensures translation of the proteins only after completion of cellularization.

As for any biological mechanism that employs probability, the question of the precision and robustness of the outcome arises. The wedge shape mesoderm folding relies on relative differences in contractility between the cells, not on absolute levels, such that the system is less restricted. Furthermore, the connectivity between cells, mediated by cell junctions, may average out local perturbations. The tension along the anterior-posterior axis of the embryo was shown to be greater than that along the DV axis, leading to the distinct elongated shape of mesoderm invagination (Chanet et al., 2017). This global tension may also act as a buffering element for local perturbations.

### Timing of actomyosin constriction by intron delay

Incorporation of timing to the interpretation of positional information adds another layer of complexity. For example, in the case of outgrowth and patterning of the vertebrate limb undergoing growth in parallel to patterning. In such systems, growth may take place following an earlier patterning phase, within a field comprised of a small number of cells. Alternatively, growth may coincide and act as an active component of patterning, whereby the more distal cells that appear at later times are relatively distant from the morphogen source, and hence are exposed to lower morphogen levels (Wolpert, 1969).

Timing is also a central issue when coordinating the processes of cellularization and mesoderm invagination. The generation of the mRNA gradient for the T48 and Mist proteins regulating actomyosin constriction takes place at the early to middle stages of NC 14. Within 10-15 minutes the Dorsal-nuclear gradient is stabilized (Rahimi et al., 2019), and in the following 15 minutes the mRNA gradient is produced. However, the process of cellularization, that is driven by expansion of the cell membranes towards the basal side, is completed only 30-40 minutes after the onset of NC 14. It is therefore imperative to assure that Myosin II recruitment driving apical constriction of the gastrulating cells, will take place only after the completion of cellularization (Dawes-Hoang et al., 2005), as premature apical constriction may be deleterious.

Most early zygotic genes, including the ones expressed in the future mesoderm, are intron-less or exhibit introns of only a few kb (Artieri & Fraser, 2014). The *T48* and *fog* genes stand out, displaying introns of 25 kb. This large intron size is conserved in the homologous genes in different *Drosophila* species. Since the speed of Pol II progression is ∼2kb/min (Fukaya et al., 2017), and is roughly similar for all genes, these introns could provide a delay of 10-15 minutes in the production of the mature mRNA. We have demonstrated that this delay indeed takes place, by comparing the detection timing of 5’ vs. 3’ *T48* probes, and by demonstrating that an mRNA produced by an intron-less *T48* gene will mature before the a transcript produced from the endogenous locus (Figure 7). It is interesting that the intron size of *T48* and *fog* are similar, facilitating a coordinated time of mature mRNA appearance. The T48 protein may generate a pattern, while the production of Fog, a diffusible extracellular ligand, may provide a temporal switch. If the Fog receptor, Mist, is produced in a graded manner, it will contribute to the graded activation. Since the *mist* gene is only 9.3 kb long, Mist protein may appear before the presentation of Fog. The manifestation of intron delay in timing has been previously demonstrated for the *kni related* gene in the *Drosophila* embryo (Rothe et al., 1992), and for *her7/hes7* in the segmentation clock in zebrafish and mice (Lewis, 2003; Takashima et al., 2011). Since the speed of transcription is uniform, intron length provides an effective temporal design mechanism in situations where intricate control of timing is necessary (Swinburne & Silver, 2008).

In conclusion, the Dorsal-nuclear gradient is utilized not only for subdivision of the embryo into distinct domains, but also for specification of graded responses leading to tissue shape changes that culminate in gastrulation. The convergence of two distinct mechanisms for generating transcriptional gradients of genes regulating actomyosin contractility, sharpens the resulting response. Finally, temporal coordination between the graded transcriptional response and execution of cell movement, is achieved by a delayed maturation of the relevant transcripts, dictated by intron size.

## Materials and Methods

### Fly lines

The bulk of experiments were performed using wildtype (*yw*) embryos. For ectopic Toll activation, we analyzed embryos generated by females carrying *UASp-Toll*^*ΔLRR*^ and the *nos-Gal4* driver (Rahimi et al., 2016). Intronless *T48* was generated by fusing the cDNA to the *T48* mesodermal enhancer and the *eve* minimal promoter (Lim et al., 2017). The construct was generated by Genscript, cloned into pATTB and injected into attP-40 flies BestGene Inc. Movie S1 involved visualization by Lightsheet microscopy of a Myosin II (Zipper)-GFP embryo (BDSC line B-51564).

### smFISH

| Gene | Probe | # Probes | Fluorophore |
| --- | --- | --- | --- |
| *twist* | 53-2,362 | 48 | Quasar 670 |
| *T48* | 1-29,056 | 48 | QUASAR 570 |
| *T48* 5' | 1-2,491 | 48 | Quasar 570 |
| *T48* intron | 13,800-16,474 | 48 | Cal-Fluor-Red 610 |
| *T48* 3' | 27,796-20,056 | 48 | Quasar 670 |
| *mist* 5' | 6-2,524 | 48 | Quasar 570 |

Position of probes is marked from the first transcribed nucleotide. Probe sets were designed by Stellaris Probe Designer and purchased from LGC Biosearch Technologies. Three hours after egg lay (AEL) embryos were fixed for 25 min in 4% formaldehyde, washed in methanol and kept at -20°C. The following day, embryos were washed in methanol and then in ethanol, rocked in 90% xylene:10% ethanol for 1 h, post-fixed in 4% paraformaldehyde/PBS/0.05 M EGTA for 25 min, washed three times (10 min each), then incubated for 6 min with 10 µg/ml Proteinase K and post-fixed again. Embryos were transferred gradually to 10% FA (Formamide Deionized, Thermo Fisher Scientific, AM9342) in 2× SSC+10 μg/ml ssDNA preheated to 37°C and prehybridized for 30 min at 37°C. Hybridization buffer included 10% FA, 10% dextran, 2 mg/ml bovine serum albumin (Thermo Fisher, AM2616), RVC (Ribonucleoside Vanadyl Complex, NEB, S1402) and ssDNA+ tRNA in 2× SSC, containing the probe set (1 ng/μl) (Trcek et al., 2017). Hybridization was carried out overnight at 37°C. Next morning the embryos were shaken gently and incubated for another 30 min. Embryos were washed twice for 30 min each at 37°C with 10% FA in 2× SSC+10 μg/ml ssDNA and gradually transferred to PBS-0.5% Tween and mounted with Vectashield+DAPI Mounting Medium (Vector Laboratories). Fluorescence was visualized with a Zeiss LSM800 confocal microscope or Nikon Eclipse Ti2 microscope (Rahimi et al., 2019).

### Image analysis for TS detection, quantification and mRNA quantification

After acquisition, raw images were processed in MATLAB. First and for each channel, the image stack was maximally projected along the Z-axis and then the TS spots in each channel were detected similar to (Raj et al., 2008). In short, z-projected images were processed with a log-filter (size=15 px, σ=2.5 px), and the detection threshold defined as the first threshold in whose environment the number of detected TSs stays relatively constant (relative change-in log-smaller than half the maximal relative change across all thresholds-in log). Then, 8-connected pixels above the threshold were defined as distinct TSs and their intensity defined as the sum of their pixel intensities. For each embryo, the midline was then defined according to the *twi* fluorescence (or *T48* if *twi* was not available) and the distance (in pixel) between each TS and the midline calculated. To convert the pixel measurement into nuclei columns, the DAPI image was used to determine the average column distance for each embryo and all distances converted accordingly. For comparisons with Dorsal nuclear level, Dorsal levels in each nuclei column were taken from (Ambrosi et al., 2014) and rescaled so that the maximal dorsal level at the midline corresponds to 1. For each embryo, only the TSs (and mRNAs) inside a manually selected region of interest (∼400 nuclei) were analyzed. mRNA levels at each distance from the midline were calculated as the median fluorescence of all pixels with this midline distance. All Codes for TSs and mRNA analysis can be found on GitHub (https://github.com/barkailab).

### Single mRNA detection and Pol II quantification

After acquisition, raw image stacks were processed in Matlab. For each fluorescence channel, the “local” background fluorescence (median fluorescence in a moving sphere) was first determined and subtracted from the raw signal. The background-corrected image stacks were processed with a 3-dimensional log-filter, and the spots on the processed image stack detected with the mRNAcount function from (Raj et al., 2008). For each 18-connected spot (mRNA and TSs), the cumulative and maximal pixel intensity was determined from the fluorescence value of all corresponding pixels. The spots were then classified into mRNA spots (single, cytoplasmic molecules) and TS spots (multiple, transcribed mRNAs at the TS) based on their maximal intensity (see Fig 4A).

### Priming probability calculation and mathematical mRNA expression model

Based on our results, TS priming was assumed to be a stochastic, independent process and thus the primed promoter fraction (at a certain nuclei column) can be described by 1-exp(-T*p), with T being the time in NC 14 and p being the priming probability. This formula was used to calculate the activation probabilities in Figure 3E. This experimentally derived probability was used as an input for the dynamic model in Figure 5. In addition the priming probability outside the midline was assumed to be linearly related to Dorsal levels (as measured by (Ambrosi et al., 2014)) and the mRNA production rate after TS priming was assumed to either be constant (=1) between the different TSs or fixed at the midline (=1) and linearly related to the Dorsal levels at other positions. After transcription, mRNAs were assumed to be stable and thus accumulate over time.

## Acknowledgements

We thank Melanie Weilert and Julia Zeitlinger for sharing information and fruitful discussions, Shalev Itzkovitz for input and discussions and Neta Rahimi for the Zipper-GFP Lightsheet movie. The work was supported by a grant from the Weizmann Data Science Research Center to F.J., and the US-Israel Binational Science Foundation 2015063 to B-Z.S. N.B. is an incumbent of the Lorna Greenberg Scherzer Professorial chair and B-Z.S. is an incumbent of the Hilda and Cecil Lewis Professorial chair in Molecular Genetics.

---

**Figure S1.**
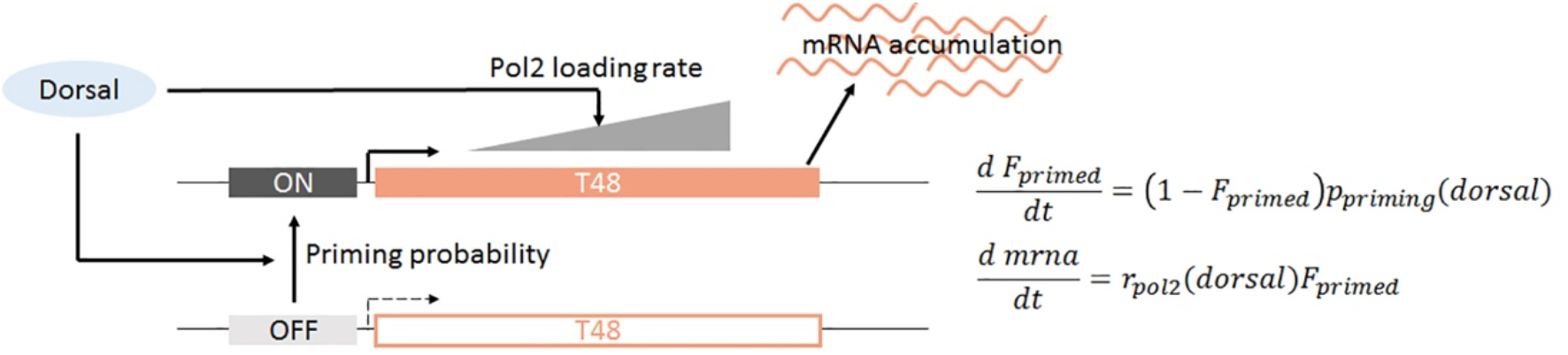
Modelling Dorsal-dependent mRNA accumulation. Based on experimental results, Dorsal was assumed to determine both the promoter priming probability and Pol II loading rate of the *T48* gene. Mature transcripts are assumed to be stable and accumulate during the course of the simulation (∼15 minutes). Differential equations used to simulate the dynamical system-Priming probability (p_priming_) and Pol II loading rate (r_pol2_) linearly depend on Dorsal level and thus on the position along the DV axis. The dynamics of the primed promoter fraction (F_primed_) and the mRNA level (mrna) is simulated for 15 minutes after model initiation (no mRNA or primed promoters at time 0). Model parameters were either arbitrary (r_pol2_ at maximal Dorsal levels) or based on experimental results (p_priming_ at maximal Dorsal=0.07 min^-1^) (Ambrosi et al., 2014).

**Figure S2.**
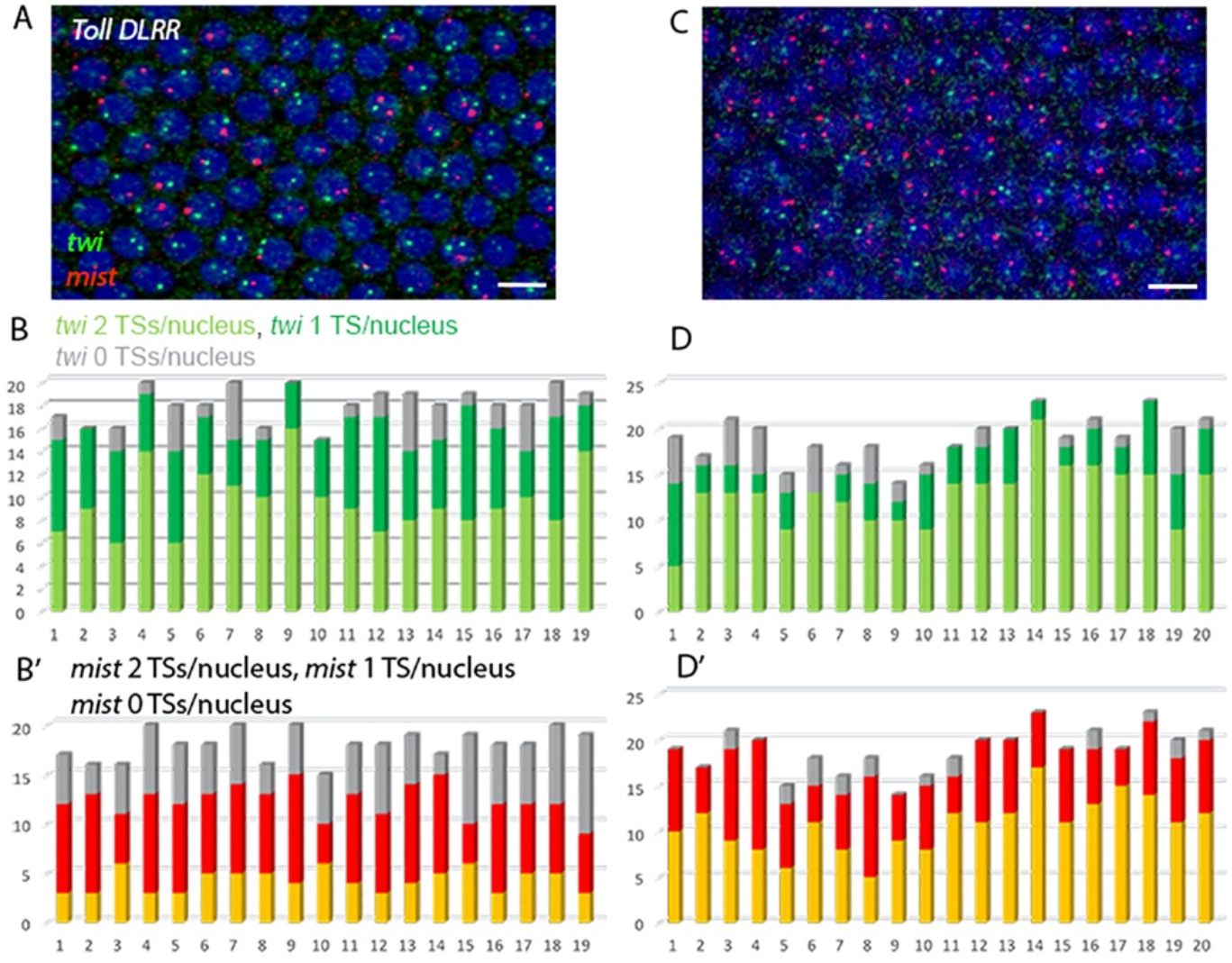
Uniform priming of *mist* in *Toll^ΔLRR^* embryos. Uniform expression of the constitutive Toll^ΔLRR^ construct drives activation of the pathway and nuclear targeting of Dorsal along the entire embryo circumference. Induction of *twi* (green) and *mist* (red) TSs responds accordingly. In a younger embryo (A-B’) one or two *twi* TSs are observed in most nuclei, while only a single or no *mist* TSs are observed in many nuclei. In an older embryo (C-D’) the majority of nuclei exhibit two TSs for both genes. Scale bar 10 μm.

## Notes

### Competing Interest Statement

The authors have declared no competing interest.

